# Widespread diversity in the transcriptomes of functionally divergent limb tendons

**DOI:** 10.1101/2019.12.18.881797

**Authors:** Nathaniel P Disser, Gregory C Ghahramani, Jacob B Swanson, Susumu Wada, Max L Chao, Scott A Rodeo, David J Oliver, Christopher L Mendias

## Abstract

Tendon is a functionally important connective tissue that transmits force between skeletal muscle and bone. Previous studies have evaluated the architectural designs and mechanical properties of different tendons throughout the body. However, less is known about the underlying transcriptional differences between tendons which may dictate their designs and properties. Therefore, our objective was to develop a comprehensive atlas of the transcriptome of limb tendons in adult mice and rats using systems biology techniques. We selected the Achilles, forepaw digit flexor, patellar, and supraspinatus tendons due to their divergent functions and high rates of injury and tendinopathies in patients. Using RNA sequencing data, we generated the Comparative Tendon Transcriptional Database (CTTDb) that identified substantial diversity in the transcriptomes of tendons both within and across species. Approximately 30% of transcripts were differentially regulated between tendons of a given species, and nearly 60% of the transcripts present in anatomically similar tendons were different between species. Many of the genes that differed between tendons and across species are important in tissue specification and limb morphogenesis, tendon cell biology and tenogenesis, growth factor signaling, and production and maintenance of the extracellular matrix. This study indicates that tendon is a surprisingly heterogenous tissue with substantial genetic variation based on anatomical location and species.

**Key Points:**

- Tendon is a hypocellular, matrix-rich tissue that has been excluded from comparative transcriptional atlases. These atlases have provided important knowledge about biological heterogeneity between tissues, and our manuscript addresses this important gap.
- We performed measures on four of the most studied tendons, the Achilles, forepaw flexor, patellar, and supraspinatus tendons of both mice and rats. These tendons are functionally distinct and are also among the most commonly injured, and therefore of important translational interest.
- Approximately one-third of the transcriptome was differentially regulated between Achilles, forepaw flexor, patellar, and supraspinatus tendons within either mice or rats. Nearly two thirds of the transcripts that are expressed in anatomically similar tendons were different between mice and rats.
- The overall findings from this study identified that although tendons across the body share a common anatomical definition based on their physical location between skeletal muscle and bone, tendon is a surprisingly genetically heterogeneous tissue.

## Introduction

Tendons are composed of a dense, hypocellular connective tissue that connect muscles to bones, and play an important role in transmitting forces throughout the musculoskeletal system during locomotion. The extracellular matrix (ECM) of tendons is composed primarily of type I collagen, in addition to other collagens, proteoglycans, and related matrix proteins (Gumucio *et al.*, 2015). Tenocytes, or tendon fibroblasts, are the most abundant cell type in tendon tissue and are responsible for ECM production, hierarchical organization, and repair (Gumucio *et al.*, 2015). Tendons are also composed of endothelial cells, sensory neurons, and tissue resident immune cells, among others, that provide a vascular supply, allow for proper motor control, and support the general growth and homeostatic functions of tendon tissue (Sugg *et al.*, 2014; Ackermann *et al.*, 2016; Noah *et al.*, 2020).

Tendons display considerable variability in architectural design and mechanical properties that are determined by articular joint geometry and locomotive functions (Biewener, 2016). Some of these functions include transmitting forces generated by large muscles capable of generating high forces, which are often referred to as prime movers of joints. These include the two major loadbearing tendons of the lower extremity ‒ the Achilles tendon, which connects the gastrocnemius and soleus muscles to the calcaneus, and the patellar tendon that attaches the quadriceps muscles to the tibia (Eng *et al.*, 2008). Other tendons are designed for fine movements and dexterity, where the muscle connected to the tendon generates less force than prime movers of the limbs and trunk, and the tendon crosses several highly mobile joints. These tendons include the digit flexors of the upper limb, which connect the flexor digitorum muscles in the forearm to the phalangeal bones in the digits (Mathewson *et al.*, 2012). Tendons may also transmit forces that help to stabilize a joint during locomotion, and help to adjust the direction of forces that prime movers exert on a joint. An example of this is the supraspinatus (SSP) tendon, which connects the SSP muscle located in the scapula to the head of the humerus, and works to guide the activity of the prime mover deltoid, pectoralis, and latissimus dorsi muscles of the glenohumeral joint (Ward *et al.*, 2006). In addition to playing an important functional role in joint movement, these four tendons are also among the most frequently injured tendons of the upper and lower extremities (Zwerver *et al.*, 2011; Colvin *et al.*, 2012; Zafonte *et al.*, 2014; Ganestam *et al.*, 2016).

The anatomical features and mechanical properties of tendons have been extensively studied (Biewener, 2016; Choi *et al.*, 2018), but less is known about underlying transcriptional differences between tendons. Comparative transcriptional atlases have provided useful knowledge about biological heterogeneity between tissues (Zhang *et al.*, 2014; Melé *et al.*, 2015; Terry *et al.*, 2018); however, tendons have been overlooked in all of the transcriptional atlases of which we are aware. Gaining greater comprehension of the transcriptional differences between tendons could help to inform our understanding of the molecular biology that specifies tendon structure and function, and also provide important insight into therapeutic interventions to treat tendon injuries and chronic degenerative conditions. Rats have been an important small animal model used to study tendon biology, but based on the growing availability of molecular genetics techniques, mice are increasingly becoming another small animal model used in basic and translational tendon research studies (Thomopoulos *et al.*, 2015). Therefore, our objective was to develop a comprehensive atlas of the transcriptome of Achilles, forepaw digit flexor, patellar, and SSP tendons of mice and rats.

## Methods

### Ethical Approval

This study was approved by the Hospital for Special Surgery/Weill Cornell Medical College/Memorial Sloan Kettering Cancer Center IACUC (protocol 2017-0035). All experiments were conducted per the approved animal protocol and were consistent with the guidelines of *The Journal of Physiology*.

### Animals

Animals used in this study were male and two months of age to be reflective of early adulthood, and due to the general similarities in overall longevity between species and growth and development milestones of animals at this time point (Mendias *et al.*, 2015; Falcioni *et al.*, 2018). C57BL/6J mice (N=4) were obtained from the Jackson Laboratory (strain 000664, Bar Harbor, ME, USA). Sprague Dawley rats (N=4) were purchased from Charles River (strain 400, Wilmington, MA, USA). A sample size of N=4 was selected based on observations of variability in transcript abundance from previous RNA sequencing (RNAseq) experiments of mice and rats (Gumucio *et al.*, 2019; Sarver *et al.*, 2019; Disser *et al.*, 2019; Talarek *et al.*, 2019). Animals were housed in specific pathogen free conditions. At the time of surgery, animals were euthanized by exposure to CO_2_ gas, followed by induction of a bilateral pneumothorax. To remove the Achilles, forepaw flexor, patellar, and supraspinatus tendons, the skin in the area was shaved, and a longitudinal incision was made through the skin that was superficial to the tendon. The surrounding connective tissue and paratenon was carefully reflected to expose the tendon. A sharp, transverse incision was made just distal to the myotendinous junction and again just proximal to the enthesis, and the tendon was carefully removed. Care was taken to exclude any non-tendon tissue in the harvest. The procedure was performed bilaterally, and tendons from each limb were combined into a single sample.

### RNA Isolation

RNA was isolated as modified from previous studies (Gumucio *et al.*, 2014; Grinstein *et al.*, 2018; Disser *et al.*, 2019). Tendons were finely minced and then pulse homogenized in Qiazol (Qiagen, Germantown, MD, USA) containing 1μg of glycogen (Qiagen) using a TissueRuptor (Qiagen). RNA was isolated with an miRNA Micro Kit (Qiagen) supplemented with DNase I (Qiagen), and quality was assessed using a BioAnalyzer RNA Pico kit (Agilent Technologies, Santa Clara, CA, USA).

### RNA Sequencing

RNAseq was performed by the Weill Cornell Epigenomics Core using a HiSeq 4000 system with 50 bp single end reads (Illumina, San Diego, CA, USA). To make libraries for RNAseq, full length double stranded cDNA (dscDNA) was generated using the SMART-Seq version 4 Ultra Low Input kit (Takara Bio USA, Mountain View, CA) and the Nextera XT DNA Library Preparation kit (Illumina). Briefly, 10ng of RNA were used to obtain first strand cDNA using SMART-Seq2 template switching and extension with SMARTScribe reverse transcriptase. cDNA was amplified using 9 cycle of PCR with SeqAmp DNA polymerase. The resulting dscDNA was validated by determining size (∼1kb) using a BioAnalyzer DNA High Sensitivity kit (Agilent Technologies). Then, 500pg of the dscDNA underwent tagmentation to generate fragments of ∼350bp containing adapter sequences. Unique indexes for each library were added by PCR amplification, and libraries were pooled together for sequencing. The pools were clustered on a single read flow cell and sequenced for 50 cycles on a HiSeq 4000 sequencer (Illumina). CASAVA (v2.20, Illumina) software was used to perform image capture, base calling and demultiplexing. Read quality was assessed and adapters trimmed using fastp (Chen *et al.*, 2018). Reads were then mapped to either the mouse genome version mm10 (UCSC, Santa Cruz, CA, USA) or rat genome version rn6 (release 95, Ensembl, Cambridge, UK) using HISAT2 (Kim *et al.*, 2019). For comparison of mouse and rat datasets, we used the getLDS function of biomaRt (Durinck *et al.*, 2009) to find homologous annotated features between the two species. Using this approach, 19,036 homologous pairs were found, but 12,555 mouse annotated features did not have rat homologs in the biomaRt database. In order to identify any additional homologous genes not identified by biomaRt, we first found homologous annotated features by peptide sequence homology using BLAST (> 90% query coverage and E-value < 1.0e-10). An additional 592 genes were identified as likely homologs using peptide similarity. Finally, for mouse annotated features for which no peptide sequence was readily available, we used nucleotide sequence to BLAST for similar annotated features in rat (> 90% query coverage and E-value < 1.0E-10). Nucleotide BLAST search identified an additional 916 annotated features as likely homologs to produce a final set of 20,544 homologous annotated features between mice and rats.

Differential gene expression analysis was calculated using edgeR (Robinson *et al.*, 2010). Genes with low expression levels (< 3 counts per million in at least one group) were filtered from all downstream analyses. A Benjamini-Hochberg false discovery rate (FDR) procedure was used to correct for multiple observations, and FDR adjusted q values less than 0.05 were considered significant. Using a power of 0.80, a *post hoc* analysis was performed (Hart *et al.*, 2013) to determine log_2_ fold change cut-offs for transcriptome comparisons between tendons and species. Sequence data is available from NIH GEO (ascension number GSE138541). Gene enrichment analysis was performed using Ingenuity Pathway Analysis (Qiagen, Valencia, CA, USA). The online Comparative Tendon Transcriptional Database (CTTDb) was built using the Data Reporting and Mining Analytics (DRaMA) platform based on R-Shiny, and contains interactive modules for data visualization, quantification, and gene enrichment analysis.

## Results and Discussion

To comprehensively evaluate the transcriptome of different limb tendons, we performed RNAseq of the Achilles, forepaw digit flexor, patellar, and SSP tendons of mice and rats. The Comparative Tendon Transcriptional Database (CTTDb), which was generated from this sequencing data, is available at https://mendiaslab.shinyapps.io/ComparativeTendonAtlas/. Approximately 150 million reads were generated for mouse tendons and 100 million reads for rat tendons (Figure 1A). Pearson correlation matrices between tendons are shown in Figure 1B. The groups that displayed the highest correlation were rat Achilles and rat patellar tendons (R^2^=0.922) and mouse Achilles and mouse forepaw flexor tendons (R^2^=0.849), while the groups that had the lowest correlation were mouse patellar and rat SSP tendons (R^2^=0.179) and mouse Achilles and rat SSP tendons (R^2^=0.190). Principal component (PC) analysis was then performed to assess variance across tendon samples. General similarities in variance were observed between Achilles and patellar tendons, and between flexor and SSP tendons (Figure 1C). We then conducted a detailed analysis of transcriptional changes, first by focusing on differences between tendons within a species (Figures 2-4, Table 1) and then comparing individual tendons across species (Table 2 and Figure 5).

**Table 1.** Gene enrichment analysis of tendons within a species. P-values of select differentially regulated pathways from mouse and rat Achilles, forepaw digit flexors, patellar, and supraspinatus (SSP) tendons that were identified using Ingenuity Pathway Analysis. ND, not significantly different.

| Pathway or Cellular Process | Achilles<br>vs Flexor | Achilles<br>vs<br>Patellar | Achilles<br>vs SSP | Flexor vs<br>Patellar | Flexor vs<br>SSP | Patellar<br>vs SSP |
| --- | --- | --- | --- | --- | --- | --- |
| *Mouse* |  |  |  |  |  |  |
| Actin Cytoskeleton Signaling | 1.26E-03 | 6.31E-12 | 4.07E-10 | 2.51E-14 | 9.33E-06 | 1.32E-09 |
| AMPK Signaling | 2.09E-08 | 7.41E-08 | 1.29E-07 | 2.00E-11 | 3.02E-06 | 1.12E-08 |
| Axonal Guidance Signaling | 3.31E-10 | 8.91E-09 | 2.51E-22 | 7.94E-17 | 5.01E-15 | 7.94E-16 |
| BMP signaling pathway | 4.27E-06 | 1.74E-02 | 1.95E-08 | 1.91E-07 | 4.90E-06 | 1.26E-05 |
| Calcium Signaling | 4.57E-07 | 3.98E-13 | 3.47E-10 | 8.32E-06 | 1.05E-06 | 4.47E-09 |
| Epithelial Adherens Junction<br>Signaling | 4.27E-09 | 7.41E-08 | 1.00E-14 | 1.05E-10 | 5.89E-06 | 1.15E-08 |
| ERK/MAPK Signaling | 4.90E-07 | 3.98E-06 | 5.62E-06 | 2.51E-11 | 5.13E-09 | 7.94E-08 |
| Fatty Acid β-oxidation I | 2.45E-05 | 1.32E-02 | ND | ND | ND | ND |
| IGF-1 Signaling | 3.72E-04 | 3.72E-03 | 1.29E-03 | 2.19E-08 | 1.00E-05 | 3.02E-04 |
| ILK Signaling | 1.55E-05 | 6.31E-12 | 1.26E-09 | 1.58E-15 | 5.62E-08 | 5.50E-10 |
| Inhibition of MMPs | ND | ND | ND | ND | ND | ND |
| p70S6K Signaling | 1.02E-04 | 6.92E-03 | 9.12E-05 | 1.91E-07 | 5.25E-05 | 9.33E-05 |
| PPARα/RXRα Activation | 1.35E-08 | 1.66E-03 | 9.77E-10 | 1.48E-08 | 1.12E-07 | 2.45E-09 |
| Protein Kinase A Signaling | 9.33E-08 | 2.63E-08 | 1.58E-16 | 4.37E-09 | 6.31E-15 | 5.01E-13 |
| STAT3 Pathway | 4.68E-06 | 5.01E-03 | 3.98E-06 | 1.58E-05 | 7.76E-07 | 1.82E-06 |
| TGFβ Signaling | 4.17E-09 | 3.89E-04 | 3.89E-10 | 2.14E-07 | 7.59E-05 | 4.79E-06 |
| Tight Junction Signaling | 2.95E-05 | 6.03E-06 | 3.47E-08 | 6.61E-07 | 8.32E-05 | 2.34E-05 |
| Wnt/β-catenin Signaling | 1.91E-09 | 2.63E-04 | 4.27E-08 | 2.45E-06 | 8.91E-06 | 7.76E-04 |
| *Rat* |  |  |  |  |  |  |
| Actin Cytoskeleton Signaling | 6.03E-06 | 7.59E-03 | 1.32E-03 | 7.08E-06 | 7.24E-04 | 1.07E-08 |
| AMPK Signaling | 3.72E-07 | ND | 9.12E-05 | 2.69E-09 | 1.32E-06 | 4.90E-06 |
| Axonal Guidance Signaling | 1.26E-17 | 7.41E-07 | 3.24E-07 | 2.00E-21 | 1.00E-18 | 1.00E-11 |
| BMP signaling pathway | 7.94E-06 | ND | ND | 5.37E-06 | 6.92E-10 | 1.32E-04 |
| Calcium Signaling | 6.17E-04 | 1.62E-03 | 6.92E-05 | 2.82E-04 | 1.95E-03 | 6.92E-07 |
| Epithelial Adherens Junction<br>Signaling | 9.55E-08 | ND | 7.08E-05 | 6.46E-08 | 9.33E-05 | 1.26E-09 |
| ERK/MAPK Signaling | 5.37E-07 | ND | 6.31E-03 | 3.98E-07 | 1.86E-04 | 3.72E-04 |
| Fatty Acid β-oxidation I | ND | ND | 2.95E-04 | 5.62E-03 | 2.51E-05 | 4.37E-04 |
| IGF-1 Signaling | 6.46E-04 | ND | 1.62E-02 | 1.29E-04 | 2.51E-06 | 9.77E-05 |
| ILK Signaling | 6.17E-08 | 2.19E-02 | 1.02E-03 | 1.23E-05 | 1.82E-05 | 2.75E-04 |
| Inhibition of MMPs | 4.68E-03 | 1.02E-07 | ND | ND | 3.09E-02 | 3.31E-02 |
| p70S6K Signaling | 5.62E-04 | ND | ND | 4.47E-05 | 6.31E-06 | 2.51E-05 |
| PPARα/RXRα Activation | 3.16E-05 | ND | 2.69E-03 | 4.90E-06 | 2.40E-08 | 9.12E-07 |
| Protein Kinase A Signaling | 1.26E-13 | ND | 8.51E-07 | 3.89E-09 | 1.55E-08 | 8.51E-08 |
| STAT3 Pathway | 3.72E-07 | ND | 4.68E-05 | 2.34E-10 | 4.68E-09 | 9.77E-08 |
| TGFβ Signaling | 1.29E-04 | ND | ND | 4.57E-05 | 2.04E-07 | 7.08E-05 |
| Tight Junction Signaling | 4.90E-07 | 2.82E-02 | 4.57E-02 | 1.07E-04 | 2.34E-02 | 1.70E-02 |
| Wnt/β-catenin Signaling | 7.94E-11 | ND | 4.90E-03 | 1.82E-08 | 7.94E-12 | 2.00E-05 |

**Table 2.** Gene enrichment analysis of tendons across a species. P-values of select differentially regulated pathways from mouse and rat Achilles, forepaw digit flexors, patellar, and supraspinatus (SSP) tendons that were identified using Ingenuity Pathway Analysis. ND, not significantly different.

| Pathway or Cellular Process | Mouse vs Rat Achilles | Mouse vs Rat Flexor | Mouse vs Rat Patellar | Mouse vs Rat Flexor |
| --- | --- | --- | --- | --- |
| Actin Cytoskeleton Signaling | 7.59E-09 | 6.03E-04 | 8.71E-10 | 3.80E-07 |
| AMPK Signaling | 2.09E-08 | 3.09E-05 | 1.95E-09 | 1.95E-06 |
| Axonal Guidance Signaling | 1.26E-12 | 2.09E-09 | 1.58E-16 | 3.98E-14 |
| BMP signaling pathway | 3.80E-02 | 5.25E-05 | 8.13E-03 | 8.91E-06 |
| Calcium Signaling | 3.72E-05 | 5.25E-03 | 3.31E-03 | 2.82E-03 |
| Epithelial Adherens Junction Signaling | 2.24E-05 | 2.29E-03 | 4.47E-06 | 4.79E-08 |
| ERK/MAPK Signaling | 4.27E-04 | 2.19E-04 | 1.91E-06 | 2.69E-05 |
| Fatty Acid β-oxidation I | 3.72E-02 | ND | ND | 1.41E-02 |
| IGF-1 Signaling | ND | 3.02E-04 | 3.80E-06 | 3.89E-04 |
| ILK Signaling | 8.71E-07 | 2.63E-04 | 5.01E-07 | 2.14E-06 |
| Inhibition of MMPs | 6.61E-04 | 3.55E-02 | ND | 2.45E-02 |
| p70S6K Signaling | 3.80E-03 | 1.55E-04 | 8.13E-06 | 2.45E-04 |
| PPARα/RXRα Activation | 2.40E-02 | 7.08E-03 | 2.34E-05 | 2.19E-03 |
| Protein Kinase A Signaling | 4.27E-09 | 1.78E-07 | 1.26E-09 | 1.10E-05 |
| STAT3 Pathway | 3.98E-04 | 4.57E-08 | 5.75E-06 | 7.24E-05 |
| TGFβ Signaling | ND | 6.61E-04 | 3.09E-03 | 2.00E-06 |
| Tight Junction Signaling | 2.14E-03 | 2.04E-05 | 8.32E-05 | 2.63E-06 |
| Wnt/β-catenin Signaling | 1.78E-02 | 7.08E-04 | 3.80E-03 | 7.59E-05 |

**Figure 1.**
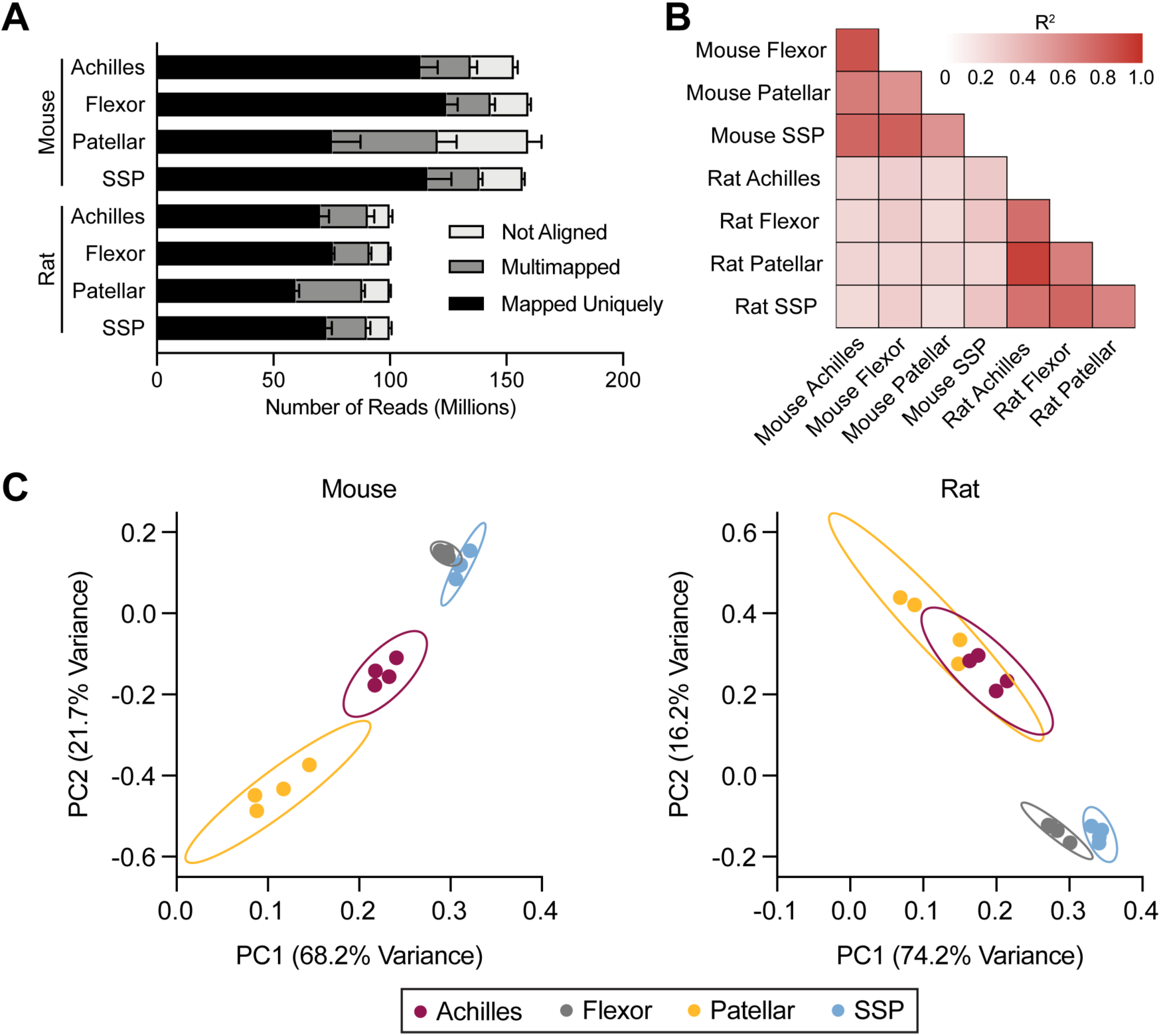
General overview. (A) Sequence alignment quantification from samples. Values are mean+standard deviation, N=4 samples per group. (B) Correlation values (R^2^) of the transcripts that are similarly expressed in tendons. (C) Principal component (PC) analysis of mouse and rat transcriptomes, with individual samples and 95% confidence interval shown.

**Figure 2.**
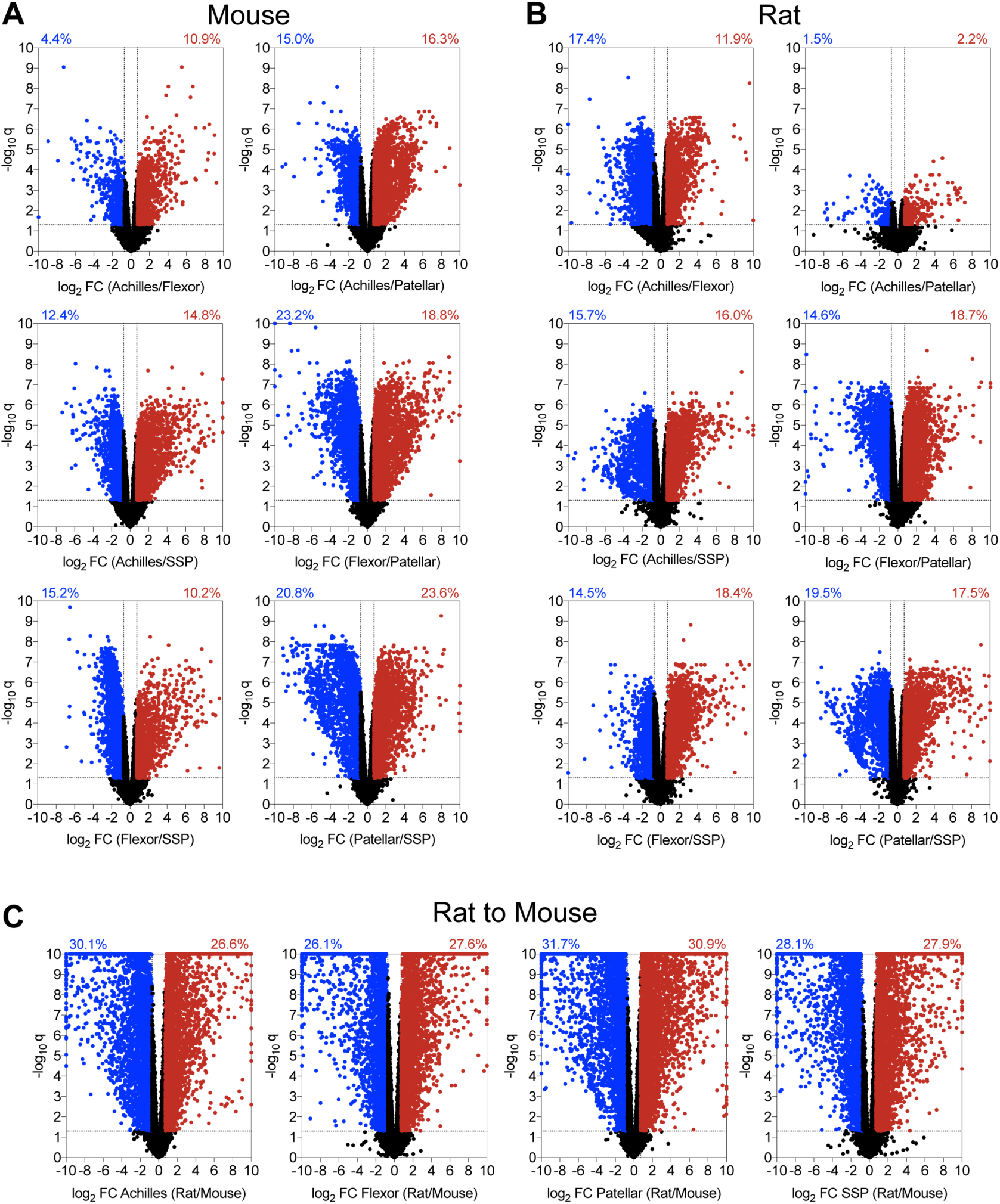
Volcano plots. Volcano plots demonstrating the log_2_ fold change and log_10_ q-values of transcripts of pairwise comparisons between tendons (A) mouse tendons, (B) rat tendons, and (C) rat to mouse tendons.

**Figure 3.**
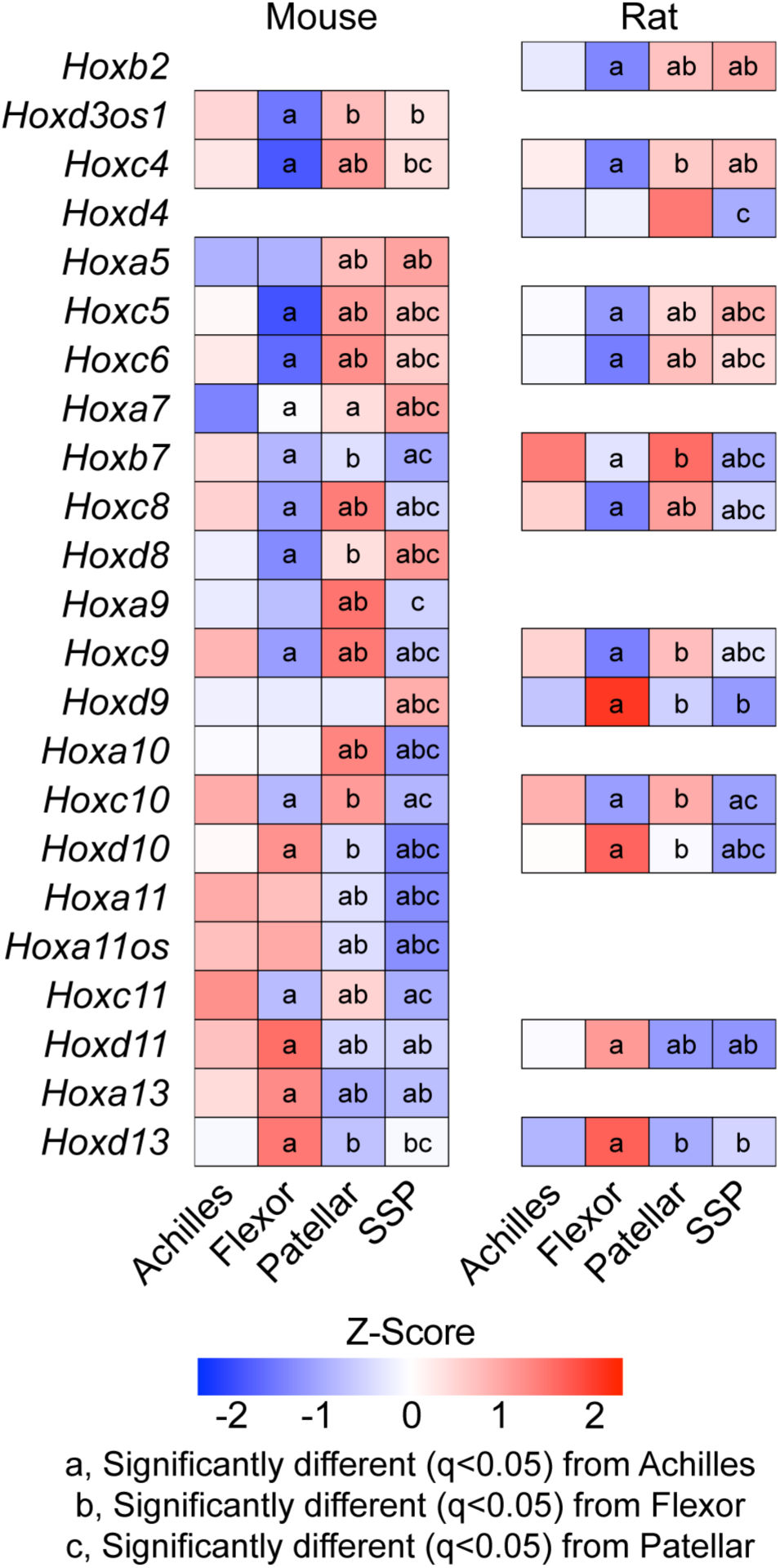
Hox genes. Heat maps of *Hox* gene expression from mouse and rat samples from Achilles, forepaw digit flexor, patellar, and supraspinatus (SSP) tendons. Values are Z-scores from the distribution of log_2_ transformed counts per million mapped reads (CPM) of mouse and rat. Differences between tendons within a species tested using edgeR: a, significantly different (q<0.05) from Achilles tendons; b, significantly different (q<0.05) from forepaw flexor tendons; c, significantly different (q<0.05) from patellar tendons.

**Figure 4.**
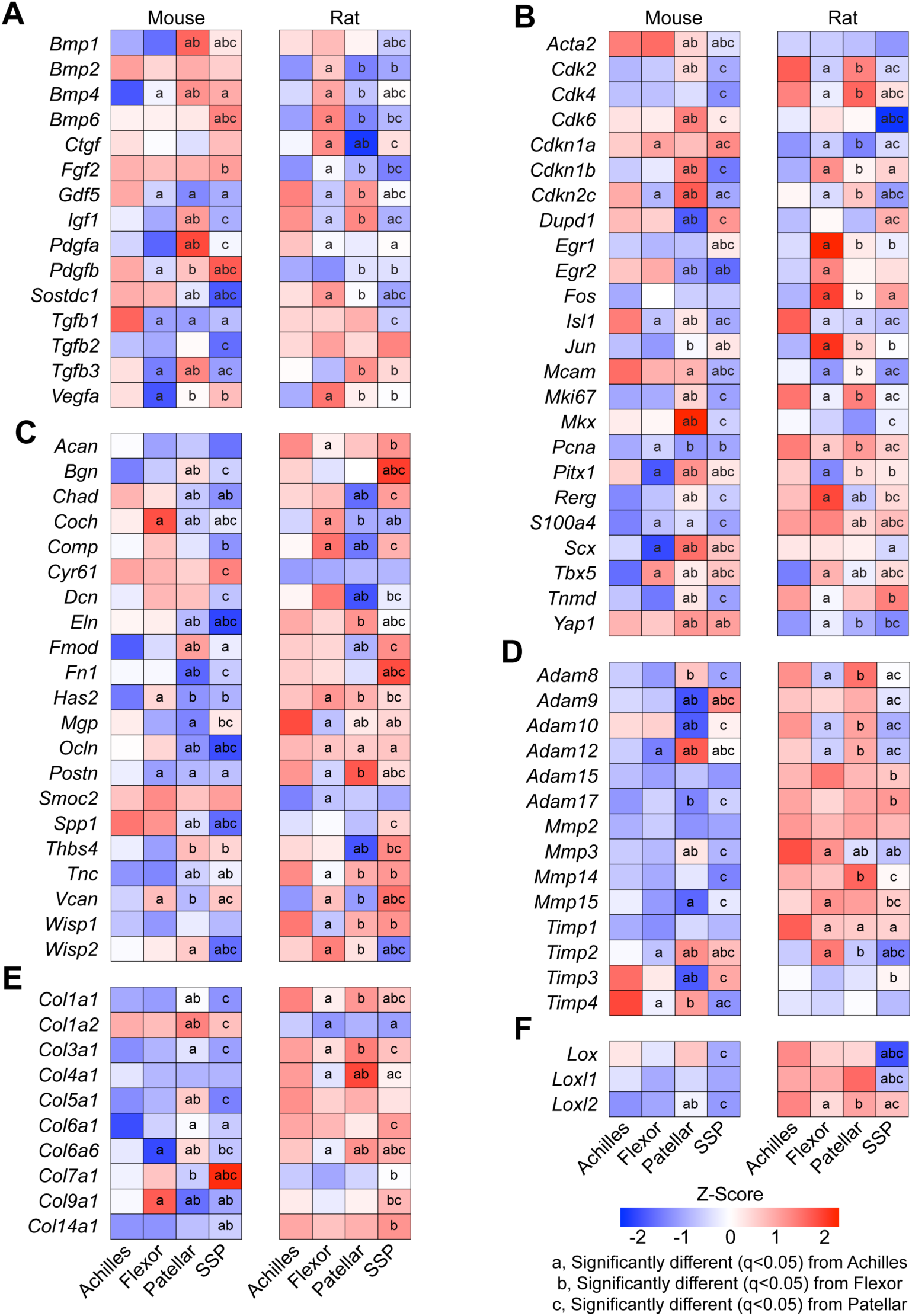
Tendon cell biology and extracellular matrix genes. Heat maps of (A) growth factor and cytokine genes, (B) genes related to fundamental cell biology or tenogenesis, (C) proteoglycans and non-collagen ECM molecules, (D) ECM proteases and their inhibitors, (E) collagens, and (F) lysyl oxidases from Achilles, forepaw digit flexor, patellar, and supraspinatus (SSP) tendons. Values are Z-scores from the distribution of log_2_ transformed counts per million mapped reads (CPM) of mouse and rat. Differences between tendons within a species tested using edgeR: a, significantly different (q<0.05) from Achilles tendons; b, significantly different (q<0.05) from forepaw flexor tendons; c, significantly different (q<0.05) from patellar tendons.

**Figure 5.**
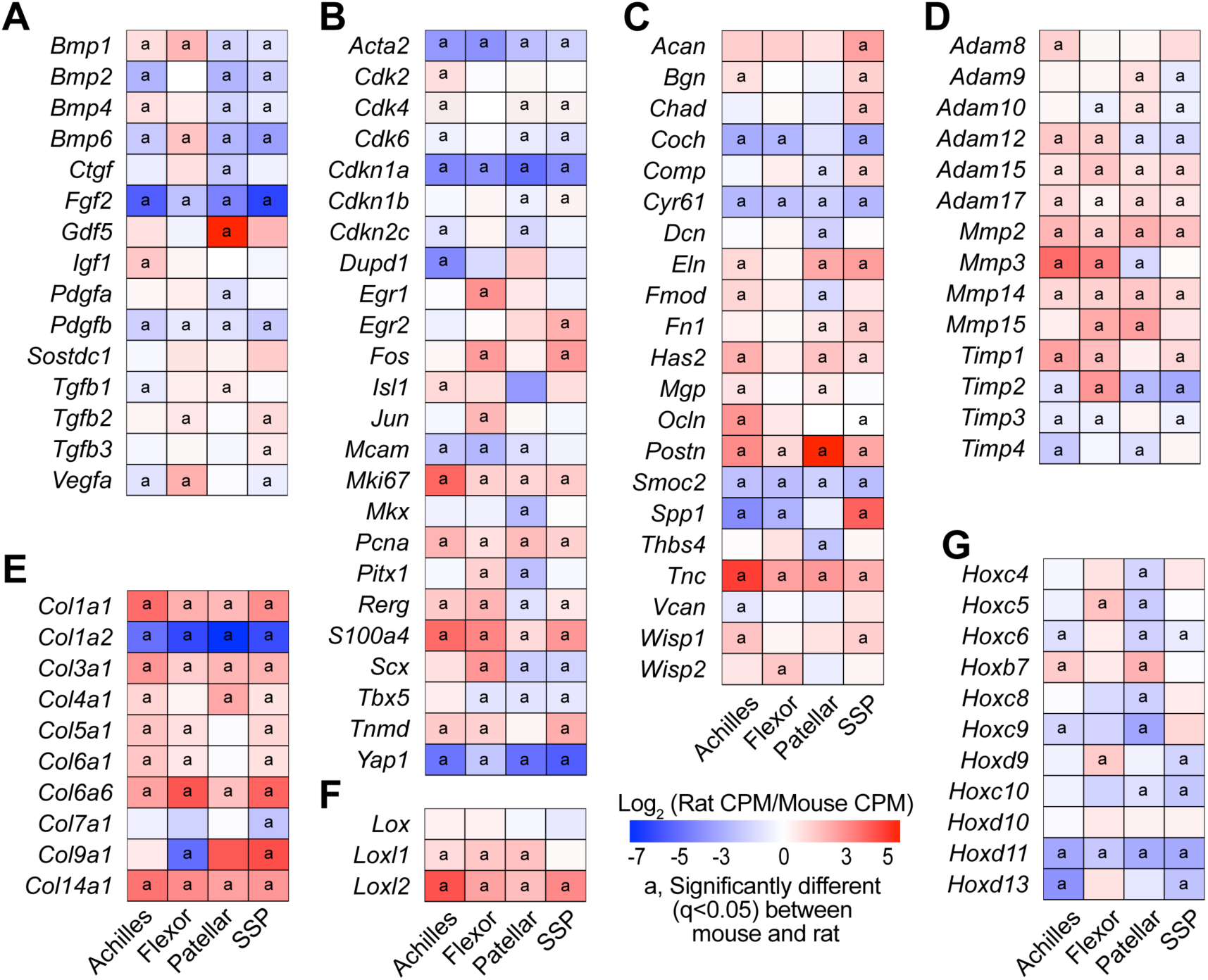
Orthologous transcriptional comparisons. Heat maps of selected genes, including (A) growth factors and cytokines, (B) genes related to fundamental cell biology or tenogenesis, (C) proteoglycans and other non-collagen ECM molecules, d(D) ECM proteases and their inhibitors. (E) collagens, (F) lysyl oxidases, and (G) *Hox* genes. Values are log_2_ counts per million mapped reads (CPM) of rat transcripts normalized to mouse transcripts. Differences between tendons within a species tested using edgeR: a, significantly different (q<0.05) from between mouse and rat samples for a given tendon.

Volcano plots were constructed to visualize general pairwise differences in the abundance and fold-change of individual transcripts between tendons within a species (Figure 2). Power analysis revealed that differences of log_2_FC of 0.72 or greater (FC = 1.65) were able to be detected with at least 0.8 power, and the threshold FC value for each volcano plot was set accordingly (Figure 2). In mice, the patellar and SSP tendons demonstrated the most divergence, with 44% of transcripts significantly differentially regulated by at least 1.65-fold (Figure 2A). Mouse flexor and patellar tendons also displayed large discrepancies, with 42% of genes significantly differentially regulated (Figure 2A). Differences in transcriptomes between the Achilles and patellar, Achilles and SSP, and the flexor and SSP tendons were 31%, 27%, and 25%, respectively (Figure 2A). The Achilles and flexor tendons demonstrated the most similarity in mice, with only 15% of transcripts significantly up or down regulated by at least 1.65-fold (Figure 2A). There were also notable discrepancies between tendon types in rats (Figure 2B). Similar to mice, the patellar and SSP tendons showed the most diversity, with 37% of genes significantly differentially regulated by at least 1.5-fold, while 33% of the transcriptome of rat flexor and patellar tendons displayed significant differences (Figure 2B). Transcripts from the flexor and SSP, Achilles and SSP, and Achilles and flexor tendons were 33%, 32%, and 29% differentially regulated, respectively (Figure 2B). Contrary to the observations made in mice, the Achilles and patellar tendons revealed the most similarity in rats, with only 4% of genes significantly differentially regulated by at least 1.65-fold (Figure 2B).

We then evaluated differences in the four tendon types across species. For Achilles tendons, 57% of genes were significantly up or down regulated by at least 1.65-fold between mice and rats, while 54%, 63%, and 56% of transcripts were similarly affected in the flexor, patellar, and SSP tendons, respectively (Figure 2C). Pathway enrichment analysis identified several pathways involved in cell signaling, growth, and ECM maintenance that were predicted to be different between mice and rats (Table 2). We then used the same gene sets that were compared between tendons within a species to analyze the same genes across species. Out of all comparisons between tendons of different species, 54% of *Hox* genes, 64% of cell proliferation and tenogenesis genes, 80% of collagen genes, 57% of proteoglycan genes, 57% of growth factor and cytokine genes, 77% of ECM metalloproteases and their inhibitors, and 58% of lysyl oxidases were differentially regulated between mice and rats (Figure 5).

The *Hox* family of transcription factors play an important role in the patterning of tissue during development, and some of the *Hox* genes continue to be expressed in tissues after development (Pineault & Wellik, 2014; Rux & Wellik, 2017). The posterior *Hox* genes (*Hox9-13*), which are involved in patterning of the limb skeleton along the proximodistal axis, were highly expressed in the four tendon types we evaluated (Figure 3, Supplemental Material S1). During development, the posterior *HoxA* and *HoxD* clusters specify both the forelimb and hindlimb, while the *HoxC* cluster is important for hindlimb tissue specification (Pineault & Wellik, 2014). Consistent with developmental expression, *Hoxc9, Hoxc10*, and *Hoxc11* display greater expression in Achilles and patellar tendons compared to the flexor and SSP tendons of adult animals (Figure 3). Furthermore, as the limb develops *Hox9* and *Hox10* paralogs are more highly expressed in the proximal region of the limb, *Hox11* is expressed in the middle regions of the limb, and *Hox13* is expressed in the distal limb regions (Pineault & Wellik, 2014; Rux & Wellik, 2017), and we observed generally similar trends in expression in many of these genes (Figure 3). *Hoxa9, Hoxc9, Hoxa10*, and *Hoxc10* were upregulated in mouse patellar tendons, which connect the proximally originating quadriceps muscle to the tibia (Figure 3). *Hoxa11* and *Hoxd11* were enriched in the Achilles and flexor tendons of mice, and *Hoxd10* and *Hoxd11* showed greater expression in the flexors of rats (Figure 3). Additionally, *Hoxa13* and *Hoxd13* in mice and *Hoxd13* in rats were abundant only in flexor tendons, which extend to the most distal regions of the limb (Figure 3). The biological function of *Hox* genes in adult tendon physiology is unknown, but as *Hox* genes are important for adult bone stem cell specification and fracture repair (Rux *et al.*, 2017; Bradaschia-Correa *et al.*, 2019) they may have a similar role in adult tendons.

We then performed pathway enrichment analysis between tendon types to examine other differentially regulated cellular functions and signaling pathways. Numerous pathways involved in growth factor signaling, cell growth and differentiation, and ECM production and degradation were identified in both mice and rats (Table 1). To further explore these processes in each tendon type, we selected panels of genes involved in these functions and report intraspecies comparisons in Figure 4 and interspecies comparisons in Figure 5.

Growth factor signaling plays an important role in regulating tendon cell biology (Sugg *et al.*, 2018; Disser *et al.*, 2019), and we observed that the expression of many growth factors differs between tendon types (Figure 4A). IGF1, which works with other growth factors to stimulate cell proliferation and protein synthesis in tendon, is essential for the proper growth of adult tendon (Disser *et al.*, 2019). Patellar tendons in mice displayed increased levels of *Igf1*, while rat patellar and Achilles tendons had higher expression of *Igf1* compared to other tendon types (Figure 4A). Expression of the cell proliferation marker *Mki67* also followed this pattern, which may indicate a role for IGF1 or other growth factors in the regulation of cell proliferation in patellar and Achilles tendons (Figure 4B). The LIM-family transcription factor *Isl1*, which is involved in transducing cytoskeletal and growth factor signaling to the nucleus (Smith *et al.*, 2014), was highly expressed in Achilles tendons of both species (Figure 4B). Other genes that encode growth factors that activate receptor tyrosine kinases, like *Ctgf* and *Fgf2*, were widely differentially regulated in rat tendons but not in mice (Figure 4A). *Gdf5* is important for articular joint formation and tendon healing (Chhabra *et al.*, 2003; Koyama *et al.*, 2008), and was expressed in much higher levels in mouse Achilles and rat Achilles and patellar tendons compared to other limb tendons (Figure 4A). Proper limb tendon development requires TGFβ signaling (Pryce *et al.*, 2009), and *Tgfb1* and *Tgfb2* displayed slight differences between tendons, with more pronounced differential regulation of *Tgfb3* (Figure 4A). Several BMP genes as well as the BMP antagonist *Sostdc1* were also differentially regulated across tendons and species, with decreased expression in patellar and SSP tendons in both species (Figure 4A). These combined results indicate extensive differential regulation of numerous growth factors and signaling molecules across tendons, which was also reflected in gene enrichment analyses.

*Scx, Mkx, Egr1*, and *Egr2* are transcription factors that play a role in tendon growth and development (Léjard *et al.*, 2011; Huang *et al.*, 2015), and were detected at varied levels depending on tendon type (Figure 4B). *Pitx1* and *Tbx5* are also transcription factors involved in limb development (Tickle, 2015) that were present at a low level in flexor and Achilles tendons, respectively (Figure 4B), which is consistent with the developmental expression patterns of these genes. Mouse patellar and rat Achilles and SSP tendons had higher expression of *Tnmd* than other tendons (Figure 4B), identifying differential regulation of this commonly used tenogenic marker (Huang *et al.*, 2015) between tendon types and species. The phosphatase *Dupd1* was expressed at very low levels in mouse and rat patellar tendons (Figure 4B, Supplemental Material S1). *Dupd1* is associated with Genitopatellar Syndrome, a rare disease that results in several physical malformations including absent patellae (Reardon, 2002; Lei *et al.*, 2016), although the specific role of DUPD1 in adult tendon homeostasis is unknown. Broadly, these differences in gene expression levels suggest potential unique mechanisms of tendon growth and development in different limb tendons.

The ECM of tendons is composed of numerous proteins that work in conjunction to transmit forces between muscles and bones. Type I collagen, which is a fibrillar molecule composed of distinct α-1 and α-2 chains, is the major protein constituent of tendons and bears much of the mechanical loads placed on tendons (Sarver *et al.*, 2017; Magnusson & Kjaer, 2019; Safa *et al.*, 2019). Patellar tendons in rats and Achilles and patellar tendons in mice demonstrated the highest level of *Col1a1* expression, but there was divergence between species for *Col1a2* expression (Figure 4E). While *Col1a1* and *Col1a2* expression in mice demonstrated nearly identical gene expression patterns, rats expressed *Col1a1* at a much higher level than *Col1a2* (Figure 4E, Supplemental Material S1). Rats also expressed much higher levels of *Col3a1*, the second most abundant fibrillar collagen in tendon, than mice (Figure 5). Several genes that encode other minor collagen proteins, such as type IV, V, VI, VII, IX, and XIV collagens were differentially regulated between tendons within a species, and most were more highly expressed in rats than in mice (Figure 5). While there was widespread differential expression in collagens between tendon types and across species, there were generally more consistent patterns of ADAM, MMP, and TIMP family expression between tendons within a species, although rats also generally displayed higher expression levels of matrix degradation enzymes (Figure 4D).

Numerous other proteins directly or indirectly interact with collagens to finely tune the mechanical properties of the tendon ECM (Thorpe *et al.*, 2013). In general, Achilles and patellar tendons are stiffer than the flexor and supraspinatus tendons (Birch, 2007; Dourte *et al.*, 2013). *Lox, Loxl1*, and *Loxl2* which encode protein crosslinking enzymes that increase matrix stiffness (Herchenhan *et al.*, 2015), as well as the matricellular protein gene *Postn* which is associated with matrix stiffness (Norris *et al.*, 2008), were elevated in rat Achilles and patellar tendons compared to supraspinatus tendons (Figure 4C-F). Proteoglycans and glycoproteins have high affinities for water, and elevated water content in tendons is associated with decreased matrix stiffness (Birch, 2007). Consistent with this, *Comp, Dcn, Fmod, Thbs4*, and *Vcan* are proteoglycan-encoding genes which were generally downregulated in Achilles and patellar tendons (Figure 4C). For ECM proteins overall (Figure 4C-F), the generally higher expression levels of collagens in rats may be due to the greater loads transmitted in rat tendons compared to mice, or reflect higher rates of matrix turnover since numerous metalloproteinases involved in collagen degradation were also elevated in rats. The differential expression of non-collagen ECM genes is likely driven by the functional demands placed upon the tissue.

There are several limitations to this study. We only evaluated male animals, as we previously observed few differences between the transcriptomes of male and female mouse Achilles tendons (Sarver *et al.*, 2017). A single time point was chosen to be reflective of early adulthood, but tendons display age related changes in structure and function (Svensson *et al.*, 2016). Additional experiments that evaluate transcriptional differences across both sexes and over the lifespan would provide important information about the heterogeneity of different tendons. We did not measure protein abundance, and it is possible that changes in the transcriptome are not reflected in the proteome. It would also be informative to determine how the transcriptomes of different tendons of mice and rats respond to bouts of mechanical loading, or in response to an injury. Despite these limitations, we feel that this study provided novel insight into basic tendon biology and will be informative in the design of future experiments.

Tendon is a structurally important connective tissue that has been largely overlooked in comparative gene expression databases. To address this, we generated an interactive database to explore the genetic differences between four commonly studied and clinically relevant tendons of mice and rats. Although we expected modest differences between tendons within a species, to our surprise we observed approximately half of the transcripts varied between some tendons within a species. This appears to be driven in part by tissue patterning signals that persist into adulthood, as well as the expression of genes that encode proteins which contribute to the unique mechanical loading needs of a specific tendon. These findings have potential clinical relevance, and suggest that factors that improve tendon healing in one tendon may not have the same effect if applied to a tendon in a different mechanical loading environment. This study also identified extensive transcriptional differences between anatomically similar tendons across species, with nearly two-thirds of transcripts displaying differential regulation. While mice and rats share many similarities in gait patterns, there are body mass distribution differences that can modulate load distribution to different limbs during locomotion (Künnecke *et al.*, 2004; Jacobs *et al.*, 2014), which likely impact the forces transmitted through anatomically similar tendons between species. This further supports the notion that the transcriptomes of tendons are finely tuned to the local biomechanical needs of the articular joints which they cross. As humans have a tendon structure and gait distinct from mice and rats, caution should be used when extrapolating findings from animal models to human tendon biology.

Future studies examining transcriptomic differences in human tendons, in combination with the CTTDb atlas, will allow results from animal models to be more effectively translated to human studies. Furthermore, gaining a greater understanding of the nature of how mechanical signals transmitted to tendons result in transcriptional changes would have important implications in the treatment of tendon injuries and diseases. In addition to mechanical signaling events, there are likely epigenetic factors that regulate gene expression, as skeletal muscles with different anatomical and functional roles display divergence in their transcriptomes and epigenomes (Terry *et al.*, 2018; Schubert *et al.*, 2019; Gumucio *et al.*, 2019). We found that even though tendons across the body share a common anatomical definition based on their physical location between skeletal muscle and bone, tendon is a surprisingly genetically heterogeneous tissue.

## Author Contributions

This study was conducted at the Hospital for Special Surgery Research Institute and the Epigenomics Core at Weill Cornell Medical College. NPD, JBS, SW, DJO, and CLM conceived and designed the study; NPD, GCG, JBS, SW, MLC, SAR, DJO, and CLM acquired, analyzed, or interpreted data; NPD, GCG, DJO, and CLM drafted the work and revised it critically for important intellectual content. All authors approved the final version of the manuscript, and agree to be accountable for all aspects of the work in ensuring that questions related to the accuracy or integrity of any part of the work are appropriately investigated and resolved

## Acknowledgements

We would like to thank Yurii Chinenov, Jonathan Daley, and Marc Sturm from the Hospital for Special Surgery for assistance in preparing the CTTDb resource.

